# The response of leaf litter bacterial communities to simulated drought depends on temperature

**DOI:** 10.64898/2026.05.05.723007

**Authors:** M. Fabiola Pulido Barriga, Claudia Weihe, Steven D. Allison, Jennifer B.H. Martiny

**Author notes:** **Address correspondence:** Jennifer B.H. Martiny, and M. Fabiola Pulido Barriga. M. Fabiola Pulido-Chavez, Department of Tropical Plant and Soil Sciences, University of Hawai’i Mānoa.

## Abstract

Microbial communities regulate carbon and nitrogen (N) cycling, yet their long-term responses to chronic global changes remain unclear. Using 12 years of grassland litter samples from the Loma Ridge Global Change Experiment in Irvine, California, we tested whether interactions between experimental drought and N deposition, and previously observed temporal variability are driven by background climatic conditions, including precipitation and temperature. Consistent with short-term studies, drought and N addition had relatively small effects on bacterial community composition compared to pronounced seasonal and interannual variability, with drought-by-year interactions explaining more variation than drought alone. Seasonal shifts were largely driven by short-term fluctuations in rainfall and temperature, whereas the substantial interannual variability in community composition was not captured by site-level climate metrics. Contrary to expectations, drought effects were influenced more by background temperature than precipitation, with the strongest effects observed in cooler years. Lastly, a bacterial taxon’s sensitivity to climate variability under ambient conditions did not predict its response to chronic drought. Together, our findings show that bacterial responses to drought are temporally dynamic and influenced by background temperature, underscoring the need for long-term longitudinal studies of soil microbial communities to better predict microbial responses under future global change.

**Importance:** Microbial responses to global change, particularly drought and nitrogen addition, are often inferred from short-term studies (< 2 years), yet natural temporal variability may overshadow experimental effects. Using a 12-year dataset of grassland leaf litter communities, we show that temporal variability, both seasonal and interannual, exert a stronger influence on bacterial community composition than chronic drought or nitrogen deposition. These findings challenge assumptions about the magnitude of drought effects, particularly in naturally drought-affected ecosystem such as California grasslands and highlight the importance of long-term datasets for predicting microbial responses to climate change. By demonstrating that bacterial communities are strongly shaped by background climatic variability (baseline precipitation and temperature independent of imposed chronic treatments) and may be buffered to sustained drought, this work improves forecasts of ecosystem responses and informs the design of global change experiments and restoration strategies in future research studies.

## Introduction

Human-driven global changes, such as drought and Nitrogen (N) deposition, are altering soil bacterial communities that drive carbon (C) and N cycling (1, 2), with particularly pronounced effects in California grasslands (3-6). Short-term drought (2–3 years) can disrupt plant-soil feedbacks (7), reduce bacterial richness and abundance and alter community composition (8, 9). Similarly, anthropogenic N deposition (10) reduces microbial biomass (11, 12) and shifts bacterial composition (13). However, long-term microbial responses to chronic drought and N deposition, as well as their potential interaction with climate variability, remain poorly understood in terrestrial ecosystems.

In ecosystems with long-term time series, mostly from marine and freshwater systems, microbial community composition is strongly influenced by natural climatic variability across seasonal and interannual timescales. For example, a five-year marine time series documented persistent annual shifts in community composition (14-16), including effects on cyanobacterial communities that lasted 2–5 years following a natural warming event (16). Similar multiyear patterns have been reported in coastal bays over seven years (17) and in freshwater lakes over six years (18). Given that terrestrial microbial communities are highly responsive to substrate moisture and temperature dynamics (19), and that marine and freshwater microbes are strongly structured by climate-linked seasonal and interannual variability (16-18), suggest that these baseline dynamics may also influence how terrestrial microbial communities responds to drought and N deposition.

Long-term microbial datasets from soil remain limited, in part due to the risk of secondary disturbance from repeated sampling. However, short-term (< 2 years) experimental drought studies in both leaf litter (20) and bulk soils (21) show that seasonal and interannual variability often explain more variation in community composition than the experimental treatments (22-24). Moreover, the magnitude of drought effects, particularly in partial rainfall exclusion experiments, depends on ambient climatic conditions (25). Consequently, accurately predicting microbial responses to global change in terrestrial ecosystems requires disentangling treatment effects from natural temporal variability and evaluating how these drivers interact.

Here, we focus on leaf litter at the soil surface (Oi horizon), where repeated sampling is possible with minimal disturbance, allowing us to track bacterial community dynamics over a 12-year period as part of the Loma Ridge Global Change Experiment (LRGCE). To simulate future global change in this region, we induced drought with rainout shelters decreasing precipitation and increased N deposition through fertilization. Short-term studies at this site showed that the treatment effects on microbial community composition are relatively small compared to large seasonal and interannual variability (13, 20). Now, with over a decade of sampling in hand, we aim to investigate whether there is a dependence between these observations. We test the following hypotheses: (1) The response of bacterial community composition to chronic changes in drought and N deposition will vary over time. (2) Ambient rainfall and temperature (i.e., “baseline” climatic variability not associated with the chronic treatments) will explain seasonal and interannual variation in bacterial community responses to the global change treatments. (3) If treatment effects vary over time, there will be no consistent correlation between a taxon’s response to background climatic variability and its response to experimental global change treatments.

## Methods

### Experimental design and sample collection

The Loma Ridge Global Change Experiment in Irvine, California (33°44′ N, 117°42′ W, 365-m elevation) was established in 2006, and the first measurement to study the effects of drought on two adjacent plant communities, a grassland and a coastal sage scrubland were taken in 2007 (26, 27). This study focuses exclusively on bacterial communities from the grassland leaf litter (Oi horizon), dominated by invasive annual grasses, including *Bromus diandrus* and *Avena fatua*. The site experiences a Mediterranean-type climate, characterized by hot, dry summers and cool, wet winters. The average annual temperature is 18.3°C, with mean lows of 15.2°C in winter and highs of 22.4°C in summer (Fig. 1). On average, ambient plots received 320 mm of precipitation annually, while drought plots, which used tarps to exclude large storms, received ∼40% less (Fig. 1). Since the beginning of the experiment, nitrogen was applied mostly as immediate-release N (5Ca(NO_3_)_2_·NH_4_NO_3_·10H_2_O), approximately twice per year, such that N plots received an average of 59 kg N ha-1 yr-1.

**Figure 1.**
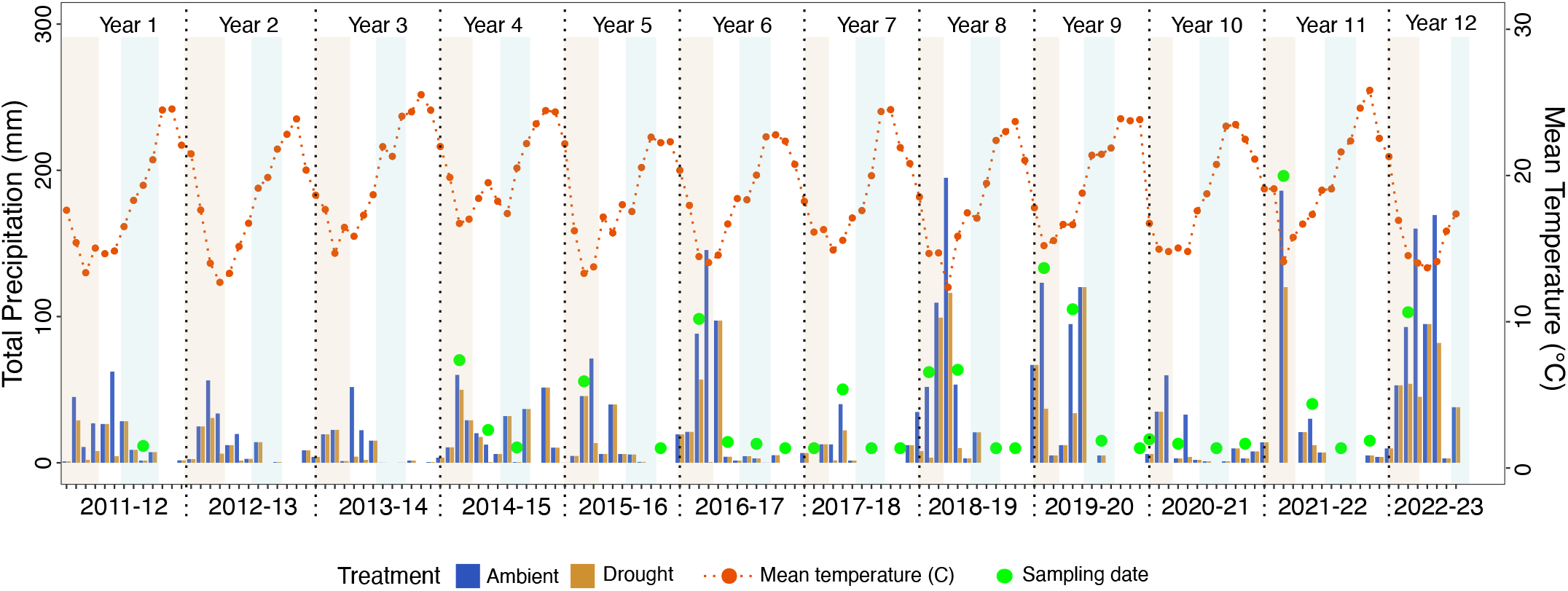
Total monthly precipitation (mm, bar plots) received per treatment (ambient vs. drought, colors) and average monthly temperature (ºC; dotted line) over a 12 year study period. In the drought plots, precipitation was reduced by approximately 40% during large rain events. Yearly time frame runs from Oct 15-Oct 14 of the following year with seasons Fall (Oct 15–Jan 14; shown in light orange), Winter (Jan 15–Apr 14), Spring (Apr 15–Jul 14; light blue), and Summer (Jul 15– Oct 14). Green dots represent the sampling dates throughout the study period. For additional details on sampling size and other metrics, see Table S1.

Recently senesced grassland litter was sampled seasonally over 12 years, starting in September of 2010. We defined seasons by grouping samples with similar precipitation and temperature conditions, using the seasonal boundaries that maximized climatic similarity among samples within each season. This approach resulted in four seasons: Fall (October 15–January 14), Winter (January 15–April 14), Spring (April 15–July 14), and Summer (July 15–October 14; Fig. 1). Additionally, “year” was defined using the California water year classification, which runs from October 1 to September 30 and encompasses one wet and one dry season (Sept-to-Sept). However, to align with our seasonal categories, we modified the water year to begin on October 15 and end on October 14 of the following year. Note, data from two years (2012-2014) are missing due to a freezer malfunction, so results presented here span the remaining 10 years of sampling (see Table S1 for additional sample size per treatment).

At each sampling timepoint, surface litter was collected from 8 replicate blocks across four treatments: ambient, drought, added N, and drought plus added N (Fig. S1). In total, plots were sampled 31 times between 2010 and 2023, resulting in 473 leaf litter samples (247 ambient, 226 drought). Litter was collected from three corners of each plot (Fig. S1). and combined in a single sterile Whirl-Pak. Samples were transported to the University of California, Irvine, where they were ground in a 70% ethanol cleaned KitchenAid coffee grinder (model BCG1110B) and stored at -80°C until DNA extraction. Additional information can be found in Table S1.

### DNA extraction and sequencing

DNA was extracted from 0.05 g of ground litter using Zymobiomics DNA extraction kits following the manufacturer’s protocol (ZymoResearch, kit D4309). The V4-V5 region of the 16S rRNA gene was amplified using the primer pair 515F and 926R (28, 29). Polymerase chain reaction (PCR) was conducted as follows: 1 µL of bacterial DNA,10.5 μl PCR grade water, 12.5 μl of AccuStartII ToughMix (2× concentration; Quantabio), and 0.5 µL each of the 10 µM barcoded forward and reverse primers. The thermocycler conditions included an initial denaturation step at 94°C for 3 min, followed by 30 cycles of 94°C for 45 s, 55°C for 30 s, 72°C for 1 min, with a final extension step at 72°C for 10 min. PCR products for each sample were pooled based on gel electrophoresis band strength and cleaned using Sera-Mag SpeedBeads (30) following previously established methods (13, 31). The pooled samples were checked for quality and quantity using the Agilent Bioanalyzer 2100. Each library (4 libraries in total) was sequenced independently using paired-end Illumina MiSeq sequencing (2 × 300 bp) at UCI’s Genomic Research Tech Hub (Irvine, CA, USA).

### Bioinformatics

Illumina data were processed using Qiime2 v2022.2.0 (32). Each sequencing run was demultiplexed and processed separately using cutadapt to remove the primers (Martin, 2011). Due to the low sequencing quality of the reverse reads, all downstream analyses were performed using only the forward reads. The forward reads were analyzed using DADA2 v2022.2 with default parameters to filter out low-quality regions, remove chimeric sequences, and generate Amplicon Sequence Variants (ASVs) (Callahan et al., 2017). DADA2 outputs from each library were combined into a single library for downstream processing, including the removal of singletons and taxonomic annotation. For taxonomic annotation, we used the SILVA v138.1 database (33) with the Qiime2 Naïve Bayes Blast+ classifier (34). To ensure that only bacterial data were analyzed, all sequences not assigned to the kingdom Bacteria or identified as mitochondria, chloroplast, unassigned and Eukaryota were removed from the final ASV table.

### Statistical analysis

The 4 Illumina MiSeq runs produced 7.2 million bacterial sequences, averaging 9,761 ASV sequences per sample, across 473 leaf litter samples. Rarefaction to 1,603 sequences per sample reduced the dataset to 414 samples (220 ambient, 194 drought) with 5,445 ASVs, which were used for downstream analyses (Table S1). Alpha diversity was assessed using the *BiodiversityR* package v2.15–4 (35). However, generalized regression models with a negative binomial distribution revealed no significant effects of drought or N on alpha diversity over time (Fig. S2). Consequently, subsequent analyses focused exclusively on bacterial community composition.

To explore drivers of litter bacterial community composition over the 12-year study period, we performed multiple Permutational Multivariate Analysis of Variance (PERMANOVA) in PRIMER v6 (36). Three sequential models were run, replacing categorical temporal variables with continuous temporal climatic components to examine how treatment effects and temporal variability are influenced by precipitation and temperature. Unless otherwise noted, all statistical analysis were performed in R (37).

In the first PEMANOVA (Original model consisting of treatment and temporal factors), we assessed the effects of categorical drought, N addition, season, and year, along with their second-order interactions, using a type III PERMANOVA with 999 permutations on square-root transformed Bray-Curtis dissimilarities. Since year significantly influenced community composition (Table S2), we further analyzed drought and N effects separately within each year. Due to the unbalanced sample sizes across years, pairwise comparisons within drought-by-year or N-by-year interactions were not feasible. Instead, we conducted independent PERMANOVA analyses for each year and treatment and visualized community composition independently for each year. To examine temporal shifts in bacterial communities, we assessed differences in community dispersion across years using the betadisper function from the vegan package and visualized mean community composition per year in NMDS space, based on the first two axes. Lastly, to identify taxa contributing to community dissimilarities across drought treatments, we performed a Similarity Percentage (SIMPER) analysis (38) on the rarefied ASV dataset. We visualized taxa contributing up to 30% of dissimilarity using ggplot2 (39).

To test whether annual variability could be attributed to precipitation and temperature patterns, we performed a second model (Annual Climate model), a Type I PERMANOVA. This model included categorical factors for season, drought, and N treatments, but we replaced “year” with total annual precipitation (mm) and mean annual maximum daily temperature (°C), and their second-order interactions. We focused on maximum daily temperatures, because temperature extremes have been shown to influence biological responses and ecosystem processes more than mean temperatures (40, 41), and because bacterial responses to climate change are expected to occur primarily through increases in seasonal temperature maxima (42).

To test whether seasonal variability in community composition was driven by seasonal precipitation and temperature patterns, we ran a third model (Seasonal Climate model), using a Type I PERMANOVA. This model included categorical year, drought, and N treatments, as well as well as continuous variables for 70-day precipitation (mm) and temperature (°C), along with their second-order interactions. The 70-day window, based on total precipitation and mean maximum daily temperature in the 70 days preceding each sampling date, was chosen to represent seasonal climatic conditions, as it effectively captured distinct seasonal patterns without overlap between sampling dates. Moreover, preliminary analyses of temporal scales (2, 14, 30, 70, and 365 days) showed that the 70-day precipitation and 2-day temperature windows explained high variance in community composition, exceeded only by the 1-year window (Fig. S3). We therefore focused on the annual and 70-day windows to identify drivers of seasonal variation. All models included block as a random effect.

Multiple analytical approaches were employed to determine whether taxa identified as drought-responsive by SIMPER, were negatively associated with precipitation and temperature, indicating a preference for drier and hotter conditions. First, we used Generalized Additive Models (GAMs) from the mgcv package (v1.9-1) to analyze the relative abundance of the top 15 ASVs, SIMPER taxa over time and across seasons. Each model included a cyclic cubic regression spline for year (k=6) to capture periodicity, a random-effect spline for ‘block’ to account for spatial variation, and a Gamma distribution with a log link. GAM model included block as random effects. Since GAMs require continuous annual sampling, we restricted our analyses to the period from 2014 to 2023 (8 years). Cyclic peaks were identified using the deriv function in the gratia package v0.9.2 to find the local maxima in the fitted splines. Second, we conducted a regression analysis with ‘plot’ as a random effect to assess the impact of cumulative annual precipitation and mean maximum daily temperature (analyzed separately) on ASV relative abundance. This approach quantified the strength and significance of each climatic variable while accounting for variability among plots. Finally, we performed Pearson correlation analysis to examine direct pairwise relationships between the top ASVs relative abundance and total annual precipitation and temperature.

## Results

The taxonomic composition of the LRGCE leaf litter communities across 12 years reflected previous studies at this site (8, 13, 31), although some taxonomic names have been updated. Pseudomonadota dominated the bacterial communities (52.3%), followed by Actinomycetota (24.6%) and Bacteroidota (20.1%). Five genera consistently dominated the communities: *Sphingomonas* (15%), *Massilia* (12.7%), *Hymenobacter* (9%), *Curtobacterium* (4.2%), and *Pseudomonas* (2.62%), collectively representing 43.6% of all bacteria across the treatments throughout the study (Fig. S4).

### The impacts of the global change treatments depend on the year

Drought and N addition treatments significantly impacted bacterial community composition, but their direct effects were minor compared to observed temporal variability in composition across seasons and years (Fig. 2a; Table S2; PEMANOVA; p < 0.05). Collectively, all factors explained 32% of the variation in community composition, with year contributing 14%, season 2%, and the year-by-season interaction, 10%. In contrast, drought and N treatments separately accounted for only 1% of the variation (p < 0.001), and the drought-by-N interaction contributed only <1% of the variation (p < 0.01; Fig. 2a; Table S2). Moreover, drought-by-year (p < 0.001) and N-by-year interactions (p = 0.05; marginally significant) explained an additional 3% and 2% of the variation, respectively (Table S2). Post hoc analyses showed significant drought effects only in four years (p < 0.05; Fig. 3). In contrast, N treatments exhibited a less consistent pattern over time, with marginally significant effects (0.05 < p < 0.1) in four of the 12 years (Fig. S5).

**Figure 2.**
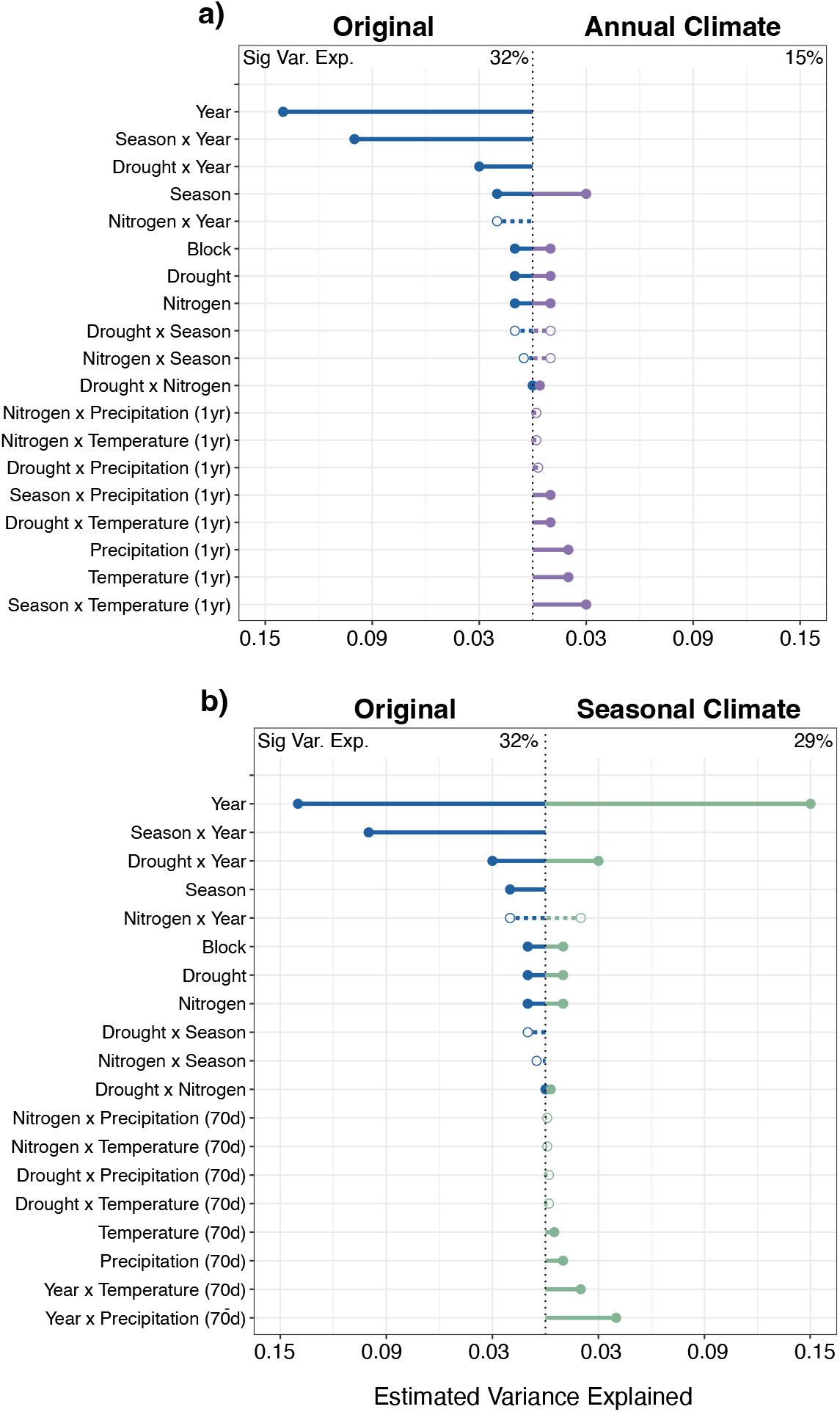
PERMANOVA analysis to test drivers of bacterial community composition for three models. **a)** The Original model tests the effects of drought, nitrogen, season, and year on bacterial community composition and the Annual Climate model replaces year with precipitation (mm) and temperature (°C) from 1 year prior to sampling. **b)** Seasonal Climate model replaces season with total precipitation (mm) and mean maximum daily temperature (°C) from 70 days prior to sampling. PERMANOVA was performed using 999 permutations on a square-root transformed Bray-Curtis dissimilarity, with block as a random effect. Significant variables (p < 0.05) are represented by solid lines and circles, whereas dashed lines with open circles are non-significant variables (p > 0.05). Total variance explained by significant variables (%) is shown at the top of the graph, with full results provided in Supplementary Table S2.

**Figure 3.**
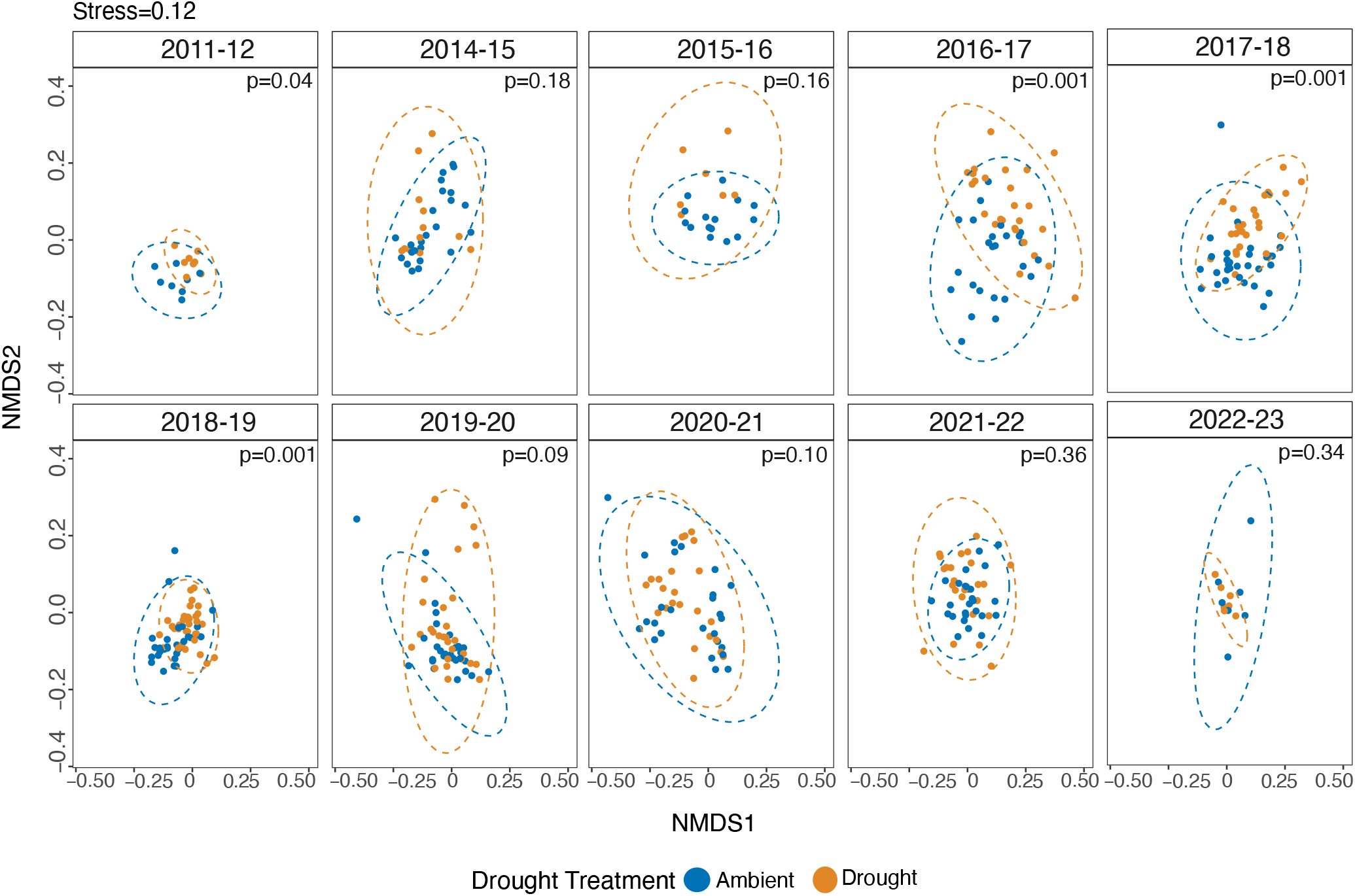
Temporal trends in bacterial community dissimilarity between ambient and drought treatments over the study period. Dissimilarity was calculated using square root-transformed Bray-Curtis distances with 999 permutations. The timeframe for each year is defined as October 15 to October 14 of the following year. Significance of differences between treatments for each year was assessed using individual PERMANOVAs with 999 permutations. The overall dissimilarity (stress) between all ambient and drought plots was 0.12 (Fig, 4). P-values represent post-hoc analysis of drought effects within each year.

### Temperature influences the response of the bacterial community to drought

We next assessed whether the observed temporal variability in bacterial community composition, including treatment-by-time interactions, could be explained by climate variability at our research site. Neither precipitation nor temperature alone fully accounted for the substantial interannual variability. Replacing the categorical “year” variable with prior-year average precipitation and mean max daily temperature in the Annual Climate model reduced the total variance explained from 32% to 15% (Fig. 2a; Table S2). Precipitation, temperature, season, and their interactions together explained only 11% of the variability in bacterial community composition, less than half of the 26% explained by the Original model (Fig. 2a; Table S2). However, annual variation in temperature, but not precipitation, partly explained the year-dependent effects of drought on bacterial community composition. Specifically, drought-by-temperature interacted to significantly affect composition (1%), whereas drought-by-precipitation did not.

An ordination plot illustrated a cyclical pattern in bacterial community structure over the 12 year study period, with bacterial communities returning to a similar composition approximately every six to seven years (Fig. 4). Bacterial community composition was also similar between years when the drought treatment had a significant impact (Years 4, 5, and 10– 12) versus years when it did not (Fig. 4). Consistent with the drought-by-temperature effect detected in the Annual Climate model, years without drought effects coincided with higher mean maximum temperatures (>23.5 °C), whereas years that drought impacted composition corresponded to lower mean annual temperature (< 23.5 °C; Fig. 4).

**Figure 4.**
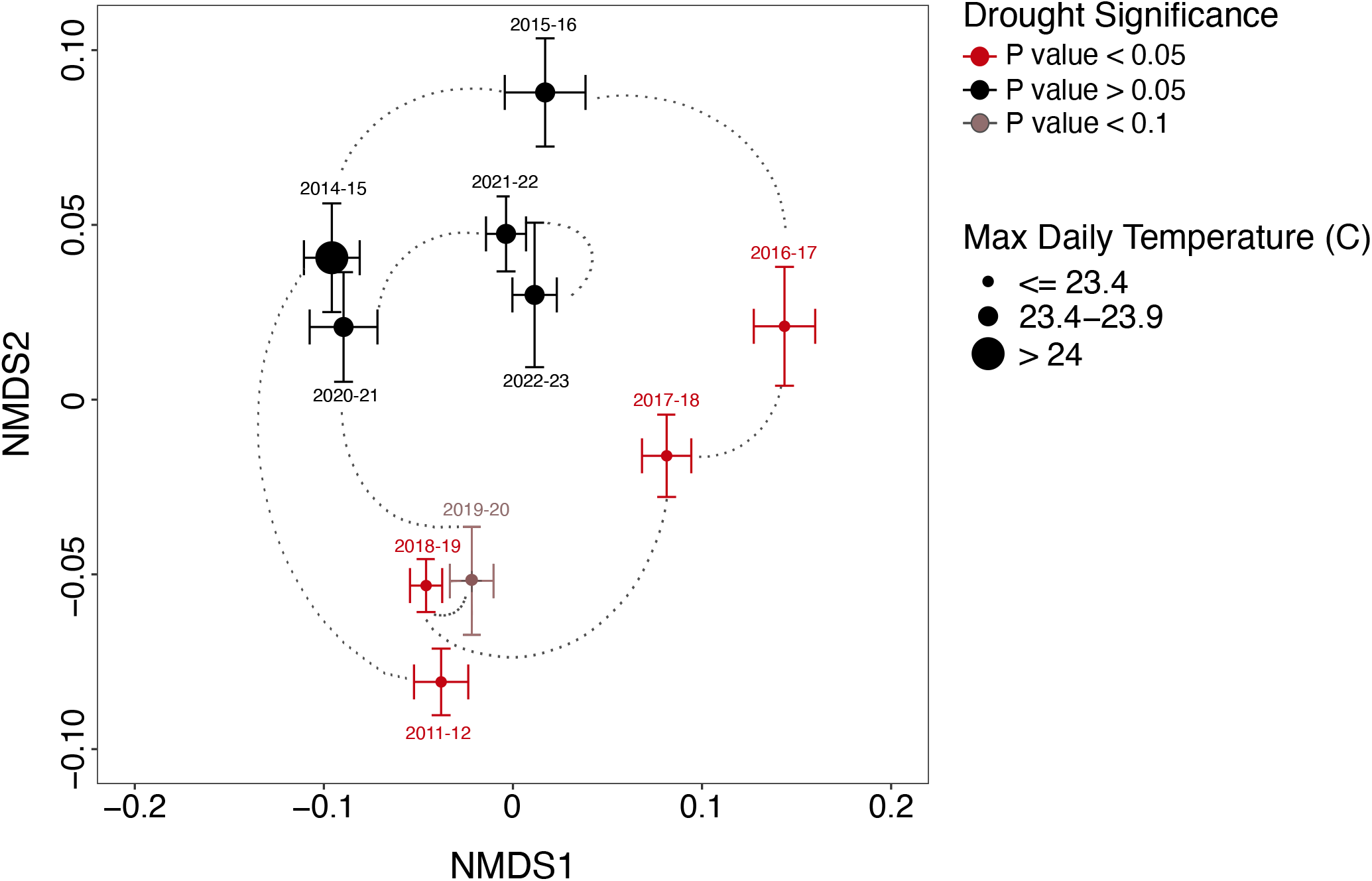
Bacterial community composition across sampling years, with dashed lines connecting the previous and subsequent years. Each symbol represents the centroid (average) community composition of all samples (across all treatments) within a sampling year with error bars showing standard error. The symbol color indicates within-year drought significance (Table 1), and symbol size represents mean maximum daily temperature (°C), with high (>23.5 °C) and low categories (<23.5 °C) defined by the dataset median as the cutoff. Drought treatment and temperature effects were categorized into two clusters and PERMANOVA showed a significant difference between non-significant (black) and significant (red) drought years (p<0.0001).

In contrast to interannual variability, precipitation and temperature over a prior 70-day window largely explained seasonal shifts in bacterial community composition (Fig. 2b; Table S2). The Seasonal Climate model accounted for 29% of community variation, closely matching the 32% explained by the Original model, which included season as a factor (Fig. 2b; Table S2). As in the Original model, interactions between treatment and 70-day precipitation and temperature were not significant in this Seasonal Climate model.

### Drought-favored taxa dominate in drier time periods

Shifts in community composition under simulated drought were primarily driven by changes in the relative abundance of 15 ASVs from 10 genera, which together explained 30% of the community dissimilarity between drought and ambient treatments (Fig. 5; Table S3). *Sphingomonas* (ASV1), *Massilia* (ASV5, ASV7), and *Curtobacterium* (ASV2) were relatively more abundant in ambient conditions, whereas *Massilia* (ASV4), *Frigobacterium* (ASV9), *Xanthomonas* (ASV11), and *Pedobacter* (ASV12) were more prevalent under drought (Fig. 5a; Table S3). These ASVs exhibited temporal fluctuations, with four of the 10 most abundant taxa displaying significant peaks in abundance 3–5 years apart (Figs. S6). Moreover, the drought-associated taxa, *Massilia* (ASV4), *Frigobacterium* (ASV9), and *Pedobacter* (ASV12) were significantly negatively correlated with precipitation, indicating a preference for drier conditions, while showing no correlation with temperature (Fig. 5b–c, Fig. S7, S8; Tables S3-S6). In contrast, ASVs that were more abundant under ambient conditions exhibited variable responses, with some taxa (e.g., *Massilia* ASV5 and *Pantoea* ASV3) increasing with temperature, while others (e.g., *Sphingomonas* ASV1, *Agrobacterium* ASV8, *Frigobacterium* ASV6, and *Sphingomonas* ASV10) decreased (Fig. 5b,c; Fig. S7,S8; Table S6).

**Figure 5.**
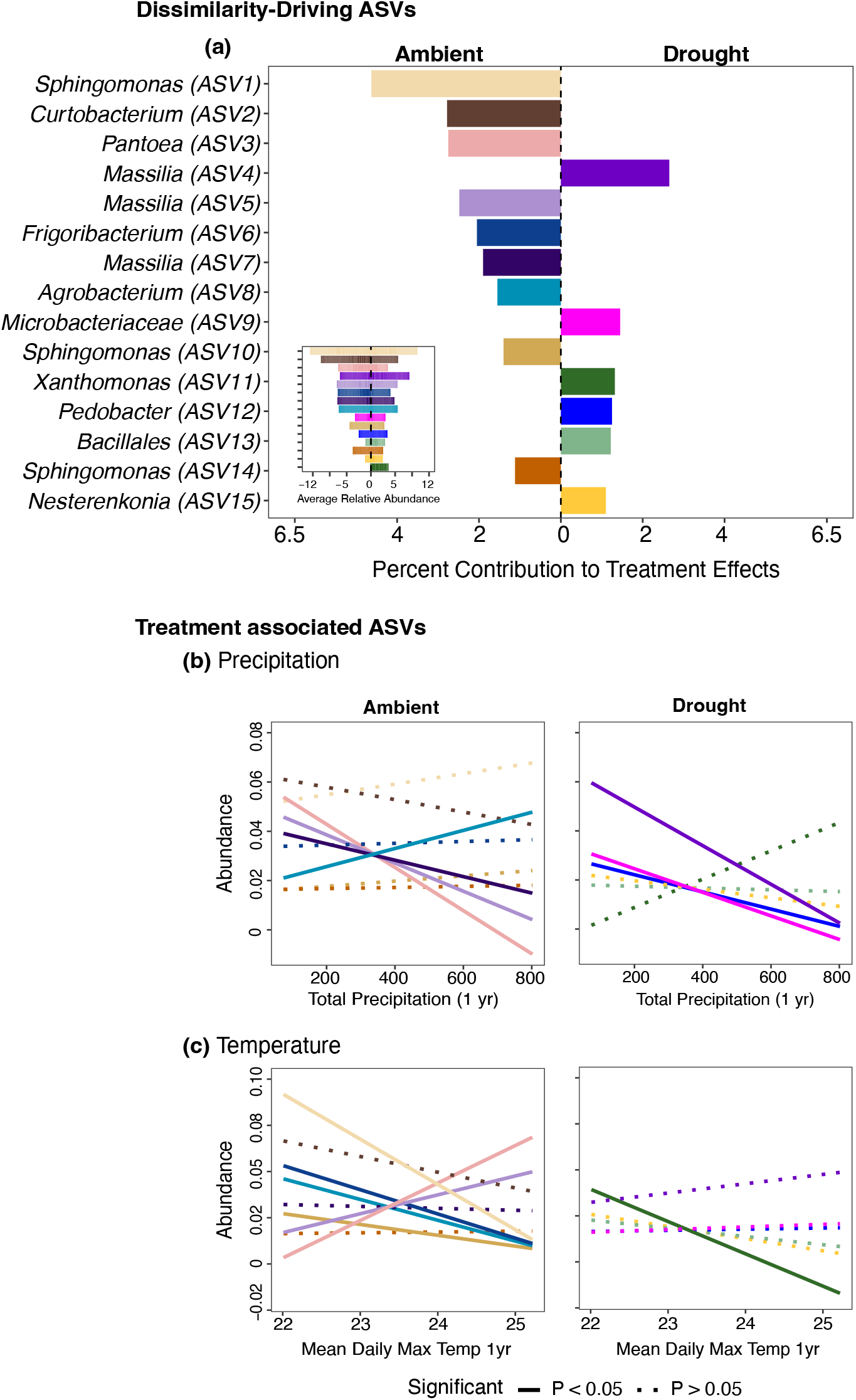
SIMPER analysis identifying (a) the top 15 ASVs across 10 genera that collectively contributed 30% of the dissimilarity between ambient and drought bacterial communities (Table S3). These taxa were detected in both treatments, but their position along the axis indicates relative abundance differences, with bars on the left showing higher abundance in ambient plots and bars on the right showing higher abundance in drought plots (see inset in panel for actual relative abundances). Panels (b, c) show correlations of ambient-versus drought-associated ASVs with (b) total annual precipitation (mm) and (c) mean maximum daily temperature (°C). Solid lines denote significant correlations (linear model and Pearson’s correlation, p < 0.05). ASVs are ranked by contribution to dissimilarity, from ASV1 (highest) to ASV15 (lowest).

## Discussion

Our long-term global change experiment shows that interannual climate variability modulates bacterial sensitivity to simulated global change treatments, such that drought, and to a lesser extent N addition, exert effects that depend on year-to-year conditions (Hypothesis 1). Over the decade analyzed, significant drought effects were observed in only five of the ten years while N addition produced marginal effects in four of ten years. This temporal contingency aligns with observations from short-term studies at this site (13, 20) and other global change experiments (21, 43), but our results extend these findings by showing that bacterial responses remain contingent over decadal timescales. Together, our findings show that bacterial community responses to drought vary greatly by year and depend on recent climate.

Because interannual variability influenced whether drought effects were significant, we sought to determine whether these temporal dynamics were driven by background climatic variability, specifically mean maximum daily temperature and mean precipitation (Hypothesis 2). While climate variables explained little of the interannual variability in bacterial composition, short-term precipitation and temperature (70 days) effectively captured seasonal dynamics. This contrast may be a response to short-term climate windows better reflecting immediate environmental conditions that affect litter associated bacteria, whereas annual averages may obscure ecologically relevant extremes such as short-term rain events or timing effects (44, 45). Moreover, categorical year may encompass processes that are not fully captured by climate metrics, including plant community dynamics (26, 46), interannual variation in litter quality and chemistry, and microbial legacy effects arising from prior disturbances or climate change history (47, 48). Indeed, drought is known to alter plant composition, with downstream effects on litter inputs and chemistry that affect bacterial community composition (49-51). At our site, a previous short-term study showed a positive correlation between plant community composition and microbial communities following the winter growing season (13), suggesting that interannual variation in plant communities may contribute to the temporal patterns observed. However, inconsistent data on plant community composition across the 12-year study period prevented us from directly testing this hypothesis.

Interannual variation in mean maximum daily temperature, but not rainfall, explained temporal variation in bacterial community responses to drought. Since the LRGCE drought manipulation (∼40% exclusion of annual precipitation) depends on background rainfall, we expected drought effects to vary with annual precipitation. Instead, drought responses were partially explained by a drought-by-temperature interaction, such that treatment effects were particularly pronounced in cooler years (<23.4 °C). In contrast, in hotter years, bacterial communities were largely unaffected by the drought treatments. Higher air temperatures increase evapotranspiration and decrease soil water potential (52). Thus, in hotter years, leaf litter bacterial communities may experience drought-like conditions regardless of background or manipulated rainfall exclusions, which could explain the lack of treatment differences between ambient and drought plots in five out of the ten years. This pattern is consistent with findings from European grasslands, where bulk surface soils (0–15 cm) with a history of drought are less sensitive to experimental drought than soils from rarely drought-exposed sites (53). It also aligns with observations that grassland plant and soil responses to warming effects are more pronounced during cooler months (54).

Although not a primary focus of this study, seasonal effects on the litter communities also varied strongly among years. This suggests that bacterial responses to seasonal shifts in temperature and precipitation may be mediated by interannual changes in litter chemistry, potentially reflecting differences in plant community inputs. This interpretation aligns other ecosystems (51, 55) and generally supports the broader idea that soil bacteria balance trade-offs between resource acquisition and stress tolerance (50, 56).

Finally, we found that sensitivity to “baseline” climate variability does not reliably predict bacterial responses to chronic drought treatment (Hypothesis 3). Although some drought-responsive taxa were more abundant during naturally dry periods (i.e., under lower precipitation or higher temperatures), these patterns were evident in both ambient and drought plots. Given that California grasslands regularly experience seasonal and multi-year droughts (3, 57, 58), it is likely that all the dominant bacterial taxa in this system are somewhat adapted to chronically low-moisture conditions, similar to native grassland plant species (59, 60). Consistent with this interpretation, *Massilia*, one of the most dominant genera in our system, has been previously detected in other drought-impacted ecosystem (61, 62), and is frequently enriched in arid or disturbed environments, including deserts, post-fire soils, and drought-adapted plant species (63-67). However, ASVs within the same genus showed contrasting responses to drought treatment. For example, both *Massilia* and another dominant genus *Curtobacterium* (which decreased under drought) possess traits associated with tolerance to environmental stress, including thermotolerance (*Massilia*), desiccation resistance (*Curtobacterium*), and lipid accumulation in both genera that stabilize cell membrane and improved water retention (63, 68-70). The presence of these stress-tolerant traits, regardless of drought treatment, suggest that they are common in bacterial taxa from drought affected ecosystem and that difference in dominance between ambient and drought plots may be due to fine-scale ecological variation among ASV rather than direct moisture or temperature responses.

## Conclusions

Our decade-long experiment revealed that the sensitivity of the leaf litter bacterial community to chronic drought is highly variable by year, indicating that short-term global change experiments may not capture microbial community responses over time. While annual climate metrics did not fully explain the variable drought response, the bacterial community responded more strongly to drought in cooler years. In contrast, short-term (70-day) climate metrics were not associated with the global change responses but did capture seasonal dynamics of bacterial community composition. The high degree of unexplained interannual variability in these soil surface communities, independent of the global change treatments, highlights the need for longer-term longitudinal studies in soil. Indeed, the relationship between climate variability and soil microbial community dynamics remains poorly characterized, let alone how future global changes will impact these dynamics.

## Acknowledgements

We thank the many members of the Martiny, Allison, and Treseder labs for their help with the maintenance and sampling of the LRGCE over the years. We thank Adam Martiny for advice on time series analyses, Alberto Barron Sandoval, Natalie Cokel, Khyati Khambel, Maanasa Ramprasad, Motahare Haghighatjoo, and Kristin Barbour for providing feedback and editing on manuscript. We also thank the Irvine Ranch Conservancy and UCI Environmental Collaboratory for facilitating research at Loma Ridge. This work was supported by the U.S. Department of Energy, Office of Science, Office of Biological and Environmental Research (grants DE-SC0016410 and DE-SC0020382) to SDA and JBHM and by the National Science Foundation (DEB-2113004) to JBHM. Finally, we thank Dr. Feizel Waffarn for his support of the UCI Chancellor’s Postdoctoral Fellowship to MFPB.

